## Supplementary Material for "Microtubule instability driven by longitudinal and lateral strain propagation"

#### This PDF includes

- Supplementary Information
- Supplementary Table 1
- Supplementary Figure 1
- Caption for Supplementary Movie 1 and Data Sets
- References

#### 1D umbrella sampling simulations

To compute the free energy distributions for the single-PF systems along the compaction reaction coordinate (RC), $\chi$, umbrella sampling simulations were carried out [1, 2]. For every nucleotide state, we first projected the stress-free simulation set $\mathbf{X}$ onto the vector $\mathbf{q}$ defining the compaction transition in the $3N$-dimensional conformational space, where $N$ is the number of atoms (the same subset of backbone atoms was used; see previous section). Here, $\mathbf{X} = (\mathbf{x}_1, \mathbf{x}_2, \ldots, \mathbf{x}_\mathrm{n})$ is a $3N \times n$-matrix of atomic positions for an ensemble consisting of $n$ structures. We then randomly drew reference configurations from the original (all-atom) trajectory such that these:

- span the full range of compaction RC values available from the stress-free simulations;
- are equidistantly spaced along the compaction RC with a step of $0.01 - 0.02$ nm;
- preferably originate from different simulation replicas;
- are separated by at least 300 ns in time, if they are drawn from the same replica.

Harmonic potentials $V_i(\mathbf{x}) = k_i[\mathbf{q} \cdot (\mathbf{x} - \mathbf{x}_i)]^2 = k_i(\chi(\mathbf{x}) - \chi_i)^2$ with spring constants $k_i = 45 - 60$ kJ mol$^{-1}$ were used to restrain the simulated structures in the proximity of the reference configurations $\mathbf{x}_i$. For technical reasons, $V_i$ were approximated with inverted flooding potentials [3]. Each of the restrained structures was simulated for at least 1 $\mu$s, and the first 100 ns were discarded. We performed 54 and 60 restrained simulations for GTP- and GDP-PF, respectively. The restrained simulations were projected onto $\mathbf{q}$ and recast in discrete form to provide unnormalized probability histograms. Following Zhu and Hummer [4], each histogram was scaled by the inefficiency factor $g_i = (1 + \tau_i)^{-1}$ where $\tau_i$ is the correlation time of projected simulation $i$. Free energy distributions were recovered using the Weighted Histogram Analysis Method [5] implemented in the BayesWHAM package [6]. Uncertainties were obtained by first sampling distribution realizations with the Metropolis-Hastings algorithm around the reconstructed distributions (see [6] for the algorithm). Then, for every point $G_l$ in the free energy distribution, Süssmann's uncertainty estimate $\delta G_l = 1/\int p_l(G)^2 dG$ was used [7, 8], where $p_l(G)$ is the distribution of free energies in bin $l$ obtained through Metropolis-Hastings sampling.

#### 2D umbrella sampling simulations

To compute the free energy landscape for the double-PF systems (Figs. 3 in the main text), 2D umbrella sampling simulations were carried out. For each nucleotide state, we first performed a 1-$\mu s$ long equilibrium simulation in the absence of axial stress ($P_{\text{zz}} = 1$ atm). We then extracted the subsets of coordinates corresponding to each of the two dimers (the same subset of backbone atoms that was used for the derivation of PF compaction), $\mathbf{X}$ and $\mathbf{Y}$, and projected those onto $\mathbf{q}$. Reference configurations from the original (all-atom) trajectory were drawn randomly such that these:

- span the full 2D area of the compaction RC plane $(\chi_x, \chi_y)$ available from the equilibrium simulation;
- are equidistantly spaced on a 2D grid with a step of $0.01 - 0.02$ nm in both dimensions;
- are separated by at least 20 ns in time.

Harmonic potentials $V_i(\mathbf{x}, \mathbf{y}) = k_i[\mathbf{q} \cdot (\mathbf{x} - \mathbf{x}_i)]^2 + k_i[\mathbf{q} \cdot (\mathbf{y} - \mathbf{y}_i)]^2$ with spring constants $k_i = 45 - 60$ kJ mol$^{-1}$ were used to restrain the simulated structures in the proximity of the reference configurations. For technical reasons, $V_i$ were approximated with a sum of two 1D inverted flooding potentials [3]. For those areas of the compaction RC plane $(\chi_x, \chi_y)$ that were not covered by the equilibrium simulation, additional reference structures were generated from neighboring umbrella windows using the structures closest to the desired value along the compaction coordinates. Each of the restrained structures was simulated for at least 500 ns and the first 100 ns were discarded. Approximately 80 restrained simulations sufficed to recover the free energy landscape in the desired area of the compaction RC plane, yielding a total of $2 \times 80 \times 0.5$ $\mu$s $= 80$ $\mu$s of sampling time for both nucleotide states. Free energy landscapes and their uncertainties (Fig. S1) were calculated as described in the previous section.

#### Calculation of the relative lateral bond stability $\Delta\Delta G^{\text{assoc}}$

According to the definition given in the main text, the relative lateral bond stability of the double-PF system is $\Delta\Delta G^{\text{assoc}} = \Delta G^{\text{assoc}}_{\text{mis}} - \Delta G^{\text{assoc}}_{\text{eq}} = \Delta G^{\text{double}}_{\text{eq}\to\text{mis}} - \Delta G^{\text{single}}_{\text{eq}\to\text{mis}}$. In general, the last two quantities are functions of the PF conformations, $\chi_1$ and $\chi_2$, and need to be obtained from the 1D and 2D free energy distributions (Fig. 2 and Fig. 3 in the main text) as follows:

$$G^{\text{double}}_{\text{eq}\to\text{mis}}(\chi_1, \chi_2) - \Delta G^{\text{single}}_{\text{eq}\to\text{mis}}(\chi_1, \chi_2) = -\text{k}_\text{B}\text{T} \log \frac{p_{12}(\chi_1, \chi_2)}{p_{12}(\chi_1^{\text{ref}}, \chi_2^{\text{ref}})} + \text{k}_\text{B}\text{T} \log \frac{p_1(\chi_1)p_2(\chi_2)}{p_1(\chi_1^{\text{ref}})p_2(\chi_2^{\text{ref}})}. \quad (1)$$

As we are interested only in relative free energies, and for convenience, we set the reference compaction values such that $p_{12}(\chi_1^{\text{ref}}, \chi_2^{\text{ref}}) = p_{12}^{\text{max}}$. Choosing $\chi_1^{\text{ref}}$ and $\chi_2^{\text{ref}}$ this way guarantees that $\Delta\Delta G^{\text{assoc}}$ is always measured relative to the conformational state in which both PFs are in the expanded/compacted state when both bound to GTP/GDP.

#### Bayesian inference of the joint free energy distribution

As sufficiently accurate umbrella sampling calculations could not be performed for the three-PF systems, we approached the problem of estimating their compaction free energy landscapes using a Bayesian inference approach. We follow the Bayesian framework previously developed by A. Ferguson to reformulate and generalize the WHAM method [6]. We seek a probability distribution discretized on a grid that partitions the compaction coordinate space $\boldsymbol{\chi} = (\chi_1, \chi_2, \chi_3) \in \mathbb{R}^3$ into $M = M_1 \times M_2 \times M_3$ bins such that $\boldsymbol{\chi} \to \{\boldsymbol{\chi}_1, \ldots, \boldsymbol{\chi}_l, \ldots, \boldsymbol{\chi}_M\} \equiv \{\boldsymbol{\chi}_l\}$, where $l$ is a one-dimensional index over the $M$ bins. The sought probability distribution $p(\boldsymbol{\chi})$ can itself be discretized on the same grid such that $p(\boldsymbol{\chi}) \to \{p_1, \ldots, p_l, \ldots, p_M\} \equiv \{p_l\}$. We note that, unlike $p(\boldsymbol{\chi})$, $\{p_l\}$ is dimensionless because each value $p_l$ is scaled with the bin volume $\Delta\boldsymbol{\chi}$. We also assume that the available unbiased MD simulation data set $\mathbf{D}$ has been projected into the compaction RC space and recast in discrete form, *i.e.* $\{N_l\}$ is the unnormalized histogram over the same grid such that $\sum_{l=1}^{M} N_l = N$, where $N$ is the number of structures in $\mathbf{D}$. We then invoke Bayes' theorem:

$$P(\{p_l\}|\{N_l\}) \sim L(\{N_l\}|\{p_l\})P_0(\{p_l\}). \tag{2}$$

$P(\{p_l\}|\{N_l\})$ is the probability of an arbitrary distribution $\{p_l\}$ given the observed data $\{N_l\}$ (*posterior*), $L(\{N_l\}|\{p_l\})$ is the *likelihood* to observe the data $\{N_l\}$ for a particular choice of $\{p_l\}$, and $P_0(\{p_l\})$ is the probability to observe an arbitrary distribution $\{p_l\}$ before obtaining any data (*prior*).

The probability to obtain the unnormalized bin counts $\{N_l\}$ for a particular distribution $\{p_l\}$, $L(\{N_l\}|\{p_l\})$, is equivalent to that of throwing a $M$-sided coin $N$ times, *i.e.* it is given by the multinomial distribution:

$$L(\{N_l\}|\{p_l\}) = \frac{N!}{\prod_{l=1}^{M} N_l!} \prod_{l=1}^{M} p_l^{N_l}. \tag{3}$$

This formula follows directly from Eq. 16 in [6] if only one unbiased simulation is considered.

From a Bayesian standpoint, the prior probability is a prior belief about the shape of the sought distribution $p(\boldsymbol{\chi})$ that is then updated by the observed simulation data $\mathbf{D}$. In our particular situation, we make use of the much more accurate estimates of the probability distributions for smaller lattice subsystems (Figs. 2 and 3 in the main text) to construct a prior following a previous maximum-entropy modeling approach [9]. The idea is to obtain the least informative estimate of $p(\boldsymbol{\chi})$ by maximizing the entropy functional,

$$S[p] = -\int p(\boldsymbol{\chi})\log p(\boldsymbol{\chi})d\boldsymbol{\chi}, \tag{4}$$

while satisfying the prior knowledge

$$\begin{aligned} p_{12}(\chi_1,\chi_2) &= \int p(\boldsymbol{\chi})d\chi_3 = f_{12}(\chi_1,\chi_2), \\ p_{23}(\chi_2,\chi_3) &= \int p(\boldsymbol{\chi})d\chi_1 = f_{23}(\chi_2,\chi_3), \\ p_i(\chi_i) &= \int p(\boldsymbol{\chi})d\chi_j d\chi_k = f_i(\chi_i), \end{aligned} \tag{5}$$

where $f_{12}(\chi_1,\chi_2)$, $f_{23}(\chi_2,\chi_3)$ and $f_i(\chi_i)$ are the normalized 2D and 1D distributions for the double-PF and single-PF systems ($i = 1,2,3$). We assume that $f_{12}(\chi_1,\chi_2)$ and $f_{23}(\chi_2,\chi_3)$ are essentially the same but transposed with respect to each other such that $\chi_2$ corresponds to the same PF in each case. We further assume the identity of the distributions $f_i(\chi_i)$. This problem has a well-known analytic solution [10, 11]:

$$p^0(\boldsymbol{\chi}) = \frac{1}{\mathcal{Z}}e^{-W_{12}-W_{23}-h_1-h_2-h_3}, \tag{6}$$

where the Lagrange multipliers $W_{12}(\chi_1,\chi_2)$, $W_{23}(\chi_2,\chi_3)$, $h_1(\chi_1)$, $h_2(\chi_2)$ and $h_3(\chi_3)$ have to be tuned such that the constraints in Eq. 5 are maintained, and $\mathcal{Z}$ is the normalization constant. We emphasize the absence of the term $W_{13}$ because no prior knowledge is available about interactions between PFs that are not directly laterally coupled. This is equivalent to assuming that such non-adjacent PFs only interfere through correlations induced by short-range physical interactions $W_{12}$ and $W_{23}$.

In practice, this variational problem narrows down to optimizing $2M^2 + 3M$ values of the discretized Lagrange multipliers (a realistic value in our case is $\gtrsim$3000), which requires a robust and fast numerical solver for large-scale nonlinear optimization problems. We employed the WORHP software (version 1.12) to set up and perform the optimization of the Lagrange multipliers [12]. We set the tolerance for fulfilling the constraints to $10^{-9}$. The optimization procedure was repeated multiple times starting from random values of the Lagrange multipliers to test the robustness of the optimization. In all cases, the procedure converged to similar solutions with the average RMSD between the solutions being $\sim 10^{-7}$.

We now introduce the prior for the discrete inference problem in Eq. 2:

$$P_0(\{p_l\}) = \begin{cases} \prod_{l=1}^{M} \exp\left\{\frac{(p_l - p_l^0)^2}{2\sigma_l^2}\right\} / \sqrt{2\pi\sigma_l^2}, \text{ if } p_l^0 > 0, \\ \\ C, \text{ otherwise,} \end{cases} \tag{7}$$

where $\{p_l^0\}$ is the discretized prior, $\sigma_l$ is the uncertainty of the prior in bin $l$, and $C$ is an arbitrary constant reflecting the fact that the prior is assumed to be uniform if not accounted for by the maximum-entropy approach. The uncertainties $\sigma_l$ can be calculated by repeating the optimization in Eq. 4 while varying the constraints such that they take into account the uncertainties of $f_{12}(\chi_1, \chi_2)$, $f_{23}(\chi_2, \chi_3)$ and $f_i(\chi_i)$ known from the previous umbrella sampling simulations.

Having defined the likelihood and the prior, the inference problem transforms into exploring the posterior. For practical reasons, it is more convenient to work with the logarithm of the posterior probability,

$$\log\tilde{P}(\{p_l\}|\{N_l\}) = \sum_{l=1}^{M} N_l \log p_l + \log P_0(\{p_l\}) - \gamma(\sum_{l=1}^{M} p_l - 1), \tag{8}$$

where $\gamma$ is the Lagrange multiplier that ensures the normalization of $\{p_l\}$ and all terms that do not depend on the index $l$ are dropped.

To obtain an estimate of the full posterior distribution, we sampled locally correlated realizations of $\{p_l\}$ from the posterior using the Metropolis-Hastings scheme as described previously [6]. To satisfy the normalization constraint, we started from a normalized uniform distribution, and every proposed realization was first re-normalized before it was used to calculate the posterior probability and to generate a new proposal. To reduce the correlation between sequential samples, only every $300^{\text{th}}$ accepted sample was saved. In total, $10^9 - 10^{10}$ samples were generated from which initial samples were discarded as not belonging to the stationary distribution. Because each of the remaining samples was representative of the posterior, we calculated the mathematical expectation $\{\bar{p}_l\}$ and the standard deviation $\{\delta p_l\}$. Likewise, the corresponding joint free energy distribution was computed by first converting every $\{p_l\}$ sample into a free energy profile, $\{G_l\} = -k_{\text{B}}T \log(\{p_l\}/\Delta\boldsymbol{\chi}) + \{C_l\}$, and then calculating the mean profile $\{\bar{G}_l\}$ and the standard deviation $\{\delta G_l\}$. The arbitrary offsets $\{C_l\}$ were adjusted such that the maxima of the $\{p_l\}$ samples correspond to zero free energy.

### Supplementary Movie and Data Set captions

**Supp. Movie 1:** Animation showing the compaction transition derived from equilibrium simulations of the single-PF system in both GTP- and GDP-state (see Fig. 2 in the main text).

**Supp. Data Set 1:** (*single_PF_GTP.pdb*) Refined single-PF structure in the GTP state.

**Supp. Data Set 2:** (*single_PF_GDP.pdb*) Refined single-PF structure in the GDP state.

**Supp. Data Set 3:** (*homo_double_PF_GTP.pdb*) Refined homotypic double-PF structure in the GTP state.

**Supp. Data Set 4:** (*homo_double_PF_GDP.pdb*) Refined homotypic double-PF structure in the GDP state.

**Supp. Data Set 5:** (*three_PF_GTP.pdb*) Refined three-PF structure in the GTP state.

**Supp. Data Set 6:** (*three_PF_GDP.pdb*) Refined three-PF structure in the GDP state.

**Table 1.** Refinement statistics for the PF systems. Names of the refined systems match the Supplementary Data Set numbering.

| | **S1** | **S2** | **S3** | **S4** | **S5** | **S6** |
|---|---|---|---|---|---|---|
| **Full map resolution** (Å) | 3.5 | 3.5 | 3.5 | 3.5 | 3.5 | 3.5 |
| $\mathbf{FSC_{avg}}$ (full map) | 0.775 | 0.786 | 0.777 | 0.790 | 0.770 | 0.787 |
| **EMRinger** | 3.02 | 2.85 | 2.97 | 3.07 | 2.32 | 2.87 |
| **RMSD bonds** (Å) | 0.022 | 0.026 | 0.023 | 0.023 | 0.023 | 0.023 |
| **RMSD angles** (°) | 2.21 | 2.30 | 2.22 | 2.21 | 2.24 | 2.21 |
| **MolProbity** | 0.93 | 1.02 | 0.76 | 0.84 | 0.76 | 0.81 |
| **Clashscore** | 0.38 | 0.57 | 0.08 | 0.13 | 0.10 | 0.04 |
| **Ramachandran Favored** (%) | 95.93 | 95.99 | 96.54 | 96.18 | 96.56 | 96.38 |
| **Ramachandran Allowed** (%) | 3.89 | 3.77 | 3.40 | 3.58 | 3.40 | 3.50 |
| **Ramachandran Outliers** (%) | 0.18 | 0.24 | 0.06 | 0.24 | 0.04 | 0.12 |
| **Poor Rotamers** (%) | 1.02 | 1.17 | 0.59 | 1.10 | 0.73 | 1.17 |
| **CaBLAM flagged** (%) | 7.54 | 6.64 | 6.03 | 6.24 | 6.54 | 6.18 |

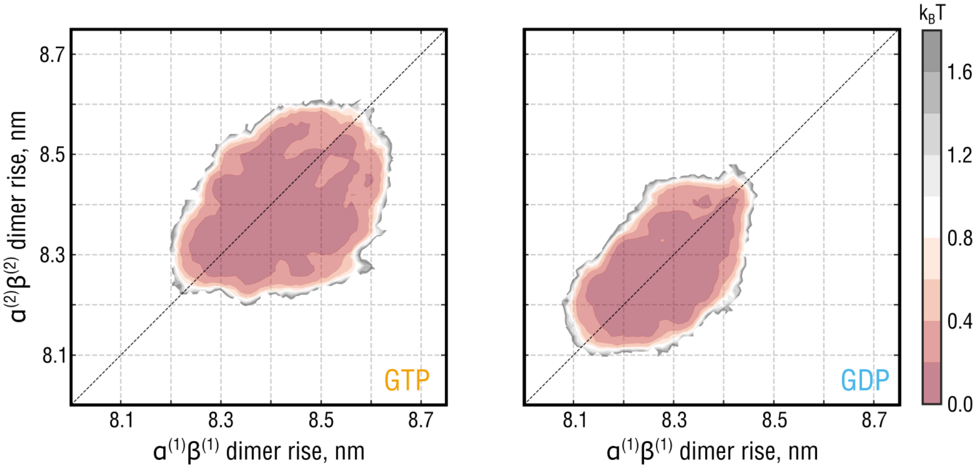


**Figure S1.** Statistical uncertainties for the 2D free energy landscapes of the system shown in Fig. 3 of the main text. Details on the error estimation are provided in the Supplementary Text.
